# The plastidial exporter Enhanced Disease Susceptibility 5 is required for the biosynthesis of *N*-hydroxy pipecolic acid

**DOI:** 10.1101/630723

**Authors:** Dmitrij Rekhter, Lennart Mohnike, Kirstin Feussner, Krzysztof Zienkiewicz, Yuelin Zhang, Ivo Feussner

## Abstract

Pipecolic acid is essential for the establishment of systemic acquired resistance in plants. It is synthesized in the plastid and further processed in the cytosol to its active form N-hydroxy pipecolic acid. Here we provide strong evidence that the exporter Enhanced Disease Susceptibility 5 is required for the biosynthesis of not only salicylic acid, but also N-hydroxy pipecolic acid, suggesting that it represents a convergent point of plant immunity.

## Introduction

Plants face numerous biotic and abiotic challenges in nature. In order to cope with these threats, they produce a variety of metabolites. These small molecules are critical for the activation of their defense system^1^. The accumulation of salicylic acid (SA) and pipecolic acid (Pip) at the site of infection as well as in systemic tissues is a key event for the successful immune response against biotrophic pathogens^2^. The first step in SA biosynthesis, the conversion of chorismic acid (CA) to isochorismic acid (IC) by Isochorismate Synthase 1 (ICS1) in plastids^3^. We previously showed that Enhanced Disease Susceptibility 5 (EDS5) is essential for the export of IC from plastids into the cytosol. IC is further processed to isochorismate-9-glutamate by avrPphB Susceptible 3 (PBS3) that subsequently decomposes to SA in a non-enzymatical process^4^. Loss of any of these three genes leads to a drastic reduction of pathogen-induced SA production and to enhanced disease susceptibility^5,6^.

Pip has been shown to be equally crucial for plant immunity as SA^7^. Its biosynthesis occurs in plastids starting from lysine^8,9^. First, the α-aminotransferase AGD2-like Defense Response Protein 1 (ALD1) catalyzes the formation of ε-amino α-keto caproic acid, which spontaneously cyclizes to Δ^1^-piperideine-2-carboxylic acid (P2C) in solution. The ketimine reductase SAR-Deficient 4 (SARD4) catalyzes subsequently the formation of Pip from P2C*^10^*. Pip-based signaling relies on Flavin-dependent Monooxygenase 1 (FMO1), which is responsible for the N-hydroxylation of Pip to yield N-hydroxy pipecolic acid (NHP)^11,12^. This newly discovered compound was proposed to be a crucial regulator of systemic acquired resistance (SAR)^11^. Although the exact subcellular localization of FMO1 has not been determined yet, studies of other FMOs strongly suggest a localization on the cytoplasmic face of the endoplasmic reticulum^13^, implying the need for a transporter from the site of Pip biosynthesis to the location where it is further processed. Both SA and NHP can be further glycosylated to form SA-β-glucoside (SAG), SA-glucoseester (SGE)^14^ and NHP-glycoside (NHP-OGlc)^12^, respectively. Glycosylation has been proposed to inactivate plant hormone signaling and is typically facilitated by cytosolic UDP-dependent glycosyltransferases (UGTs)^15^.

## Results and discussion

Not only pathogenic infection, but also abiotic stresses like ozone or UV-C treatment stimulate the biosynthesis of SA and SAG^16^, leading to similar changes in gene expression^17^. In order to have a fast and reproducible test system for Pip synthesis, we examined the possibility to induce Pip production by UV-C stress. Indeed, Pip accumulates over time in *Arabidopsis thaliana* leaves in a similar course of time (Fig. 1a), as it was described for *Pseudomonas syringae* infection before^11^. Beside Pip, we also observed increased amounts of NHP and NHP-OGlc (of which the MS/MS fragmentation patterns are depicted in Fig. S1)^12^. This indicates that Pip oxidation and glycosylation similarly occur after UV-C treatment. To exclude that the observed synthesis of Pip and NHP-OGlc is activated by UV-C-triggered SA accumulation, we examined Pip and NHP-OGlc contents in the SA-deficient mutants, *eds5, pbs3* (Fig. 1b-e) and Salicylic acid Induction Deficient 2 *(sid2),* harboring a mutation in the *ICS1* gene (Fig. S2a and b). As expected, the mutant lines, but not wild type plants, lack both SA and SAG 24 hours after being exposed to UV-C for 20 min (Fig. 1b and c). On the other hand Pip and NHP-OGlc accumulation was observed only in wild type, *pbs3* and *sid2,* but surprisingly not in *eds5* plants (Fig. 1d and e). Similar to pathogen-treated plants^11^, UV-C stress-induced Pip-biosynthesis and -processing do not depend on the presence of SA in UV-C treated plants. The absence of Pip and NHP-OGlc in *eds5* raised the question whether EDS5 is responsible for the export of not only the SA precursor IC, but also of Pip. Recently, it was shown that *fmo1* mutants are impaired in the hydroxylation of Pip to NHP. Instead, they accumulate large amounts of Pip upon UV-C irradiation (Fig. S3), again in a similar extent as described for pathogen assays^11^. In line with FMO1 functioning downstream of Pip synthesis, these results corroborate that EDS5 acts upstream of FMO1 by exporting Pip to the cytosol.

**Figure 1.**
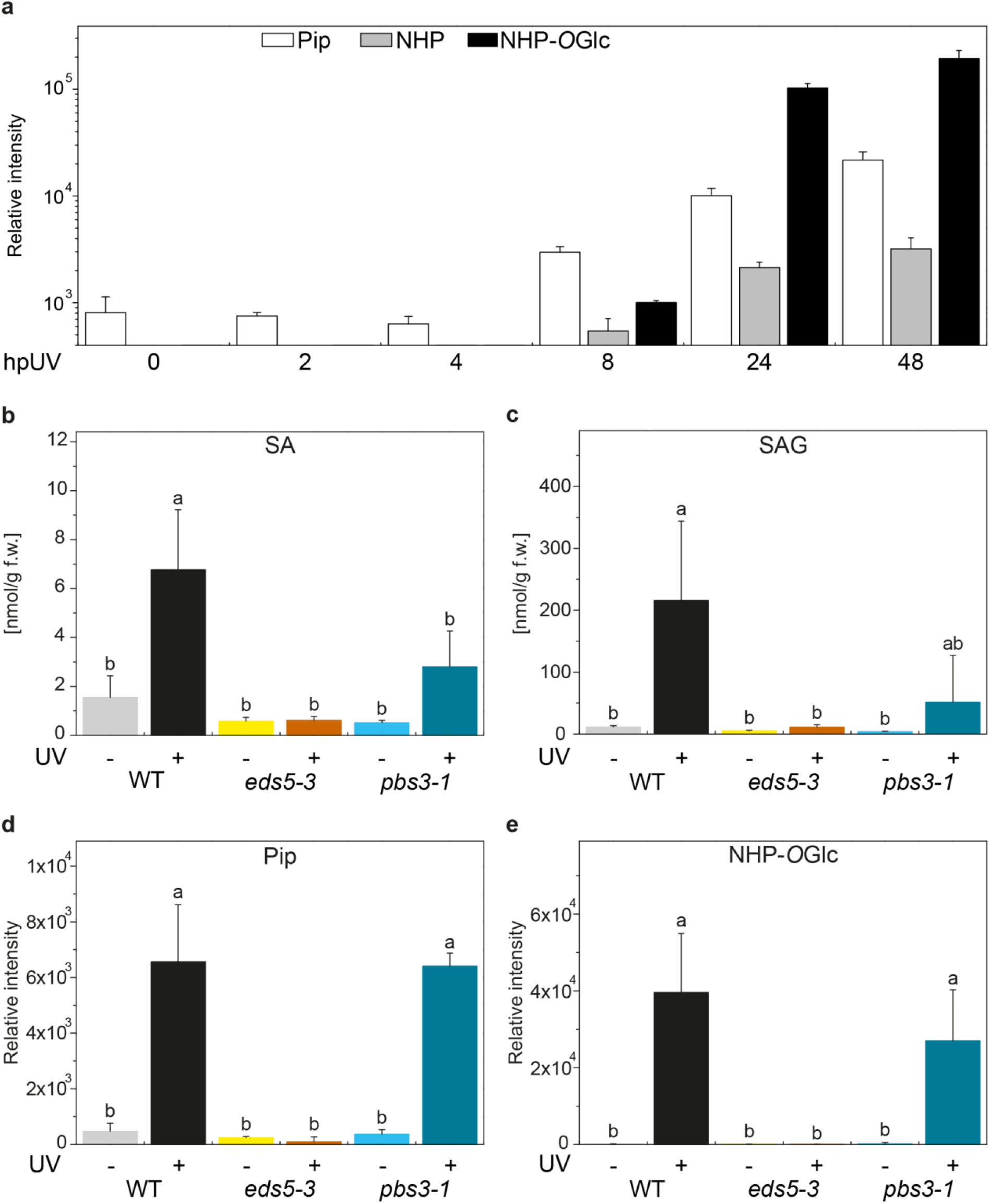
UV-C treatment of *Arabidopsis thaliana* leads to an accumulation of the signaling compounds SA and Pip and their corresponding glycosides. **a**, Levels in counts per second of Pip (white bars) and its downstream products NHP (grey bars) and NHP-OGlc (black bars) in *Arabidopsis* wild type leaves (Col-0) at different time points after UV-C light treatment. **b-e**, Levels of SA and its glycoside SAG (nmol g^-1^ leaf fresh weight [f.w.]) and levels of Pip and its glycoside NHP-OGlc (counts per second) in leaves of wild type (Col-0), respectively *eds5-3* and *pbs3-1* mutant plants 24 hours after UV-C treatment in comparison to untreated plants. Bars represent the mean ± STD of three biological replicates. Statistical differences among replicates are labeled with different letters (P < 0.05, one-way ANOVA and post hoc Tukey’s Test; n = 3). The experiments were repeated twice with similar results.

The assumed block in Pip export did not result in an enrichment of Pip in *eds5* plants, despite the induction of Pip biosynthesis on the transcriptional level: Both *SARD4* and *EDS5* were strongly upregulated upon UV-C treatment in the SA deficient *sid2* mutant and wild type plants (Fig. S2c and d), supporting our metabolite profiling results. In analogy, we observed increased levels of IC in *pbs3,* but not in *eds5* mutant plants upon UV-C treatment (Fig. S4). It is likely that a diversion of the metabolic flux prevents a harmful accumulation of both Pip and IC in plastids^18^. Pip may either inhibit its own biosynthesis via a feedback loop or feed into the lysine degradation pathway towards the Krebs cycle^19^. IC on the other hand might be channeled towards the synthesis of aromatic amino acids.

In order to exclude that the observed absence of NHP and NHP-OGlc in *eds5* mutants is due to their inability to accumulate SA, we tested whether external SA supply could induce Pip and NHP-OGlc production in *eds5* and *pbs3* plants. As reported, SA treatment triggers the biosynthesis of Pip and NHP-OGlc in wild type plants (Fig. 2a)^7^. SA biosynthesis is impaired in the *pbs3* mutant, but drenching the soil with SA still initiated Pip biosynthesis in these plants (Fig. 2a). However, as with UV-C treatment, external SA also did not lead to Pip accumulation in *eds5* mutants. This is consistent with EDS5 representing a plastidial exporter of Pip independent of SA. When Pip irrigation was used to trigger *in planta* Pip biosynthesis, we detected increased levels of the downstream product NHP-OGlc in wild type plants (Fig. 2b). In *eds5* and the Pip biosynthesis mutant *sard4,* the levels of NHP-OGlc are reduced by more than 60%. This suggests that in the mutants only the external Pip was metabolized further, whereas *de novo* Pip biosynthesis did not occur. The ability of *eds5* plants to convert exogenously applied Pip into NHP-OGlc supports FMO1 being active outside of plastids and not affected by the loss of function of *EDS5.*

**Figure 2.**
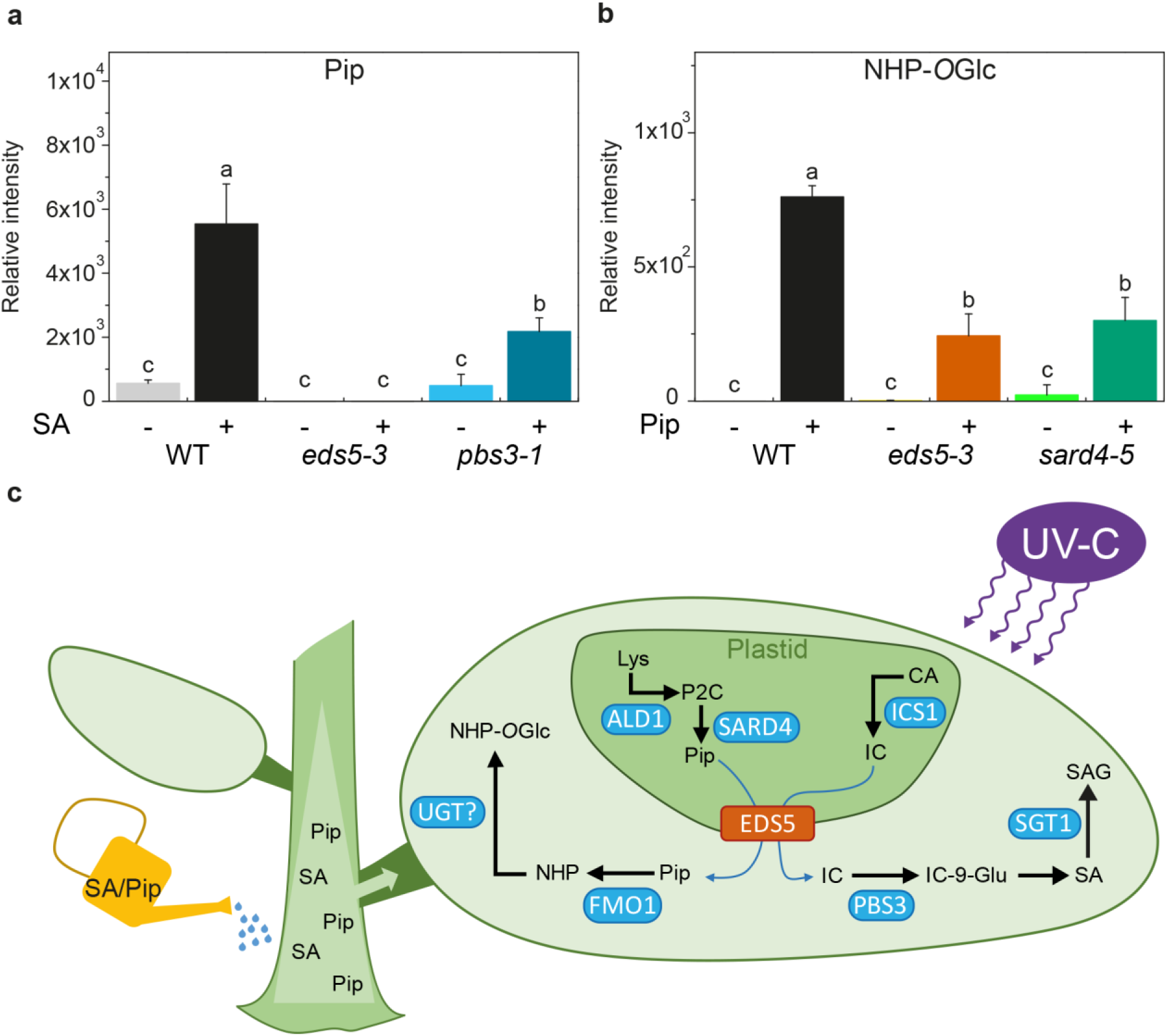
Root application of either SA or Pip induces Pip biosynthesis and processing in leaves. **a**, Levels of Pip (counts per second [cps]) in leaves of wild type (Col-0) *eds5-3* and *pbs3-1* plants 24 hours after soil drenching with water (-) or SA (+). **b**, Levels of NHP-OGlc (counts per second [cps]) in leaves of wild type (Col-0), *eds5-3* and *sard4-5* plants 24 hours after soil drenching with water (-) or Pip (+). Bars represent the mean ± STD of three biological replicates. Statistical differences among replicates are labeled with different letters (P < 0.05, one-way ANOVA and post hoc Tukey’s Test; n = 3). The experiment was repeated twice with similar results. **c**, A working model depicting EDS5 as the central hub in the biosynthesis of the signaling compounds SA and NHP. The induction of these pathways was facilitated here by UV-C light treatment.

Taken together, we show that UV-C treatment is sufficient to induce the production of Pip and its metabolites. This process does not require the presence of SA and can thus be used to study the SA-independent branch of plant defense signaling. Moreover, we identified here a previously unknown connection between Pip and *EDS5,* a gene that, so far, was only recognized for its involvement in SA biosynthesis^16^. Most likely EDS5 is also responsible for the export of Pip from plastids into the cytosol, where it is further processed by FMO1. This spatial separation of Pip biosynthesis and site of activation adds an additional layer of regulation. Our study implies that EDS5 serves as the central hub in the biosynthesis of two major defense signaling molecules, SA and NHP (Fig. 2c). It is surprising that no pathogen effector has been found to target EDS5 and thereby exploit this key point in plant immunity.

## Methods

### Plant material

*Arabidopsis* plants were grown in a chamber at 22 °C with a 16 h light period and 60% relative humidity for 4-5 weeks. For our experiments, we use *Arabidopsis* ecotype Col-0 and the following mutants in this background: *eds5-3^20^, pbs3-1^6^, sard4-5^10^*, and fmo1-1^11^, which were described previously.

### UV-C and soil drench treatment

For the UV-C treatment, we followed previous protocol^16^. In short, 4-5 week old *Arabidopsis* plants were exposed to UV-C light (254 nm) for 20 min at 50 cm distance to the lamp (TUV T8 30W, Philips) for the induction of SA and Pip biosynthesis. For the treatment with SA and Pip, we followed previously described protocols^7,20^. 4-5 week old plants, were soil drenched with either 10 mL water, 10 mL of a 5 mM Pip solution (P45850, Sigma) or 10 mL of a 5 mM SA solution (S5922, Sigma), equals to 50 μM final concentration. Samples were collected 24 hours after treatment and metabolites were extracted and analyzed as described earlier^10,21^. The deviation of exact mass to accurate mass for Pip, NHP and NHP-OGlc was less than ±2 mDa in the untargeted metabolite analysis. SA and SAG were quantified based on internal D4-SA standard (C/D/N Isotopes Inc., Pointe-Claire, Canada). The NHP standard was chemical synthesized as described in Hartmann et al., 2018^11^. The MS/MS spectra of NHP and NHP-OGlc corresponds to the results from Chen et al., 2018^12^.

### Quantitative real-time PCR analysis

To analyze the expression of *SARD4* (At5g52810) and *EDS5* (AT4G39030) after UV-C treatment, total RNA was isolated from frozen leaves with the Spectrum™ Plant Total RNA Kit (Sigma) following the manufacturer’s instruction. One microgram of RNA was treated with DNase I (Thermo Fisher Scientific) and cDNA was synthesized using Revert Aid H Minus Reverse Transcriptase (Thermo Fisher Scientific). Quantitative RT-PCR was performed using Takyon No ROX SYBR Mastermix blue dTTP (Kaneka Eurogentec) in reaction volume 20 μl. The gene ACTIN8 (At1g49240) was used as a control. Each reaction was performed with material from plants harvested from three independent samples in iQ5 real time detection system (Bio-Rad). Primers are depicted in Supplementary Table 1.

## Acknowledgments

We are grateful to Ulf Diedrichsen and Brigitte Worbs for the synthesis of the NHP authentical standard. This work was supported by the DFG (IRTG 2172 “PRoTECT” program of the Göttingen Graduate Center of Neurosciences, Biophysics, and Molecular Biosciences.) to D.R., L.M. and I.F.; I.F. was additionally supported by DFG excellence initiative (ZUK 45/2010 and INST 186/822-1). Y.Z. was supported by Natural Sciences and Engineering Research Council of Canada, Canada Foundation for Innovation, and British Columbia Knowledge Development Fund.

## Author contributions

D.R., K.F. and I.F. conceived and designed the experiments. D.R., L.M., and K.Z. performed the experiments. D.R., K.F., Y.Z., and I.F. wrote the article.

## Competing interests

Authors declare no competing interests.

## Data and materials availability

All data is available in the main text or the supplementary materials. The authors responsible for distribution of materials integral to the findings presented in this article are: Ivo Feussner and Yuelin Zhang.

## Notes

#### Summary of Updates

Title and format were updated

