## Supplemental information for "The plastidial exporter Enhanced Disease Susceptibility 5 is required for the biosynthesis of *N*-hydroxy pipecolic acid"

### Supporting information file

**Tab. S1. Primers used in this study.**

| Primer | 5'-3' sequence | Purpose |
|---|---|---|
| qPCR EDS5 fwd | ATAGACTGCTTATACCTTTCTT | expression |
| qPCR EDS5 rev | AAATCCGACGAGAACGA | expression |
| qPCR SARD4 fwd | GCGATACAGAGAGGGAGT | expression |
| qPCR SARD4 rev | GCTGAGGTAAGTCTCGTGA | expression |
| qPCR Actin8 fwd | GGTTTTCCCCAGTGTTGTTG | expression |
| qPCR Actin8 rev | CTCCATGTCATCCCAGTTGC | expression |

### Figures

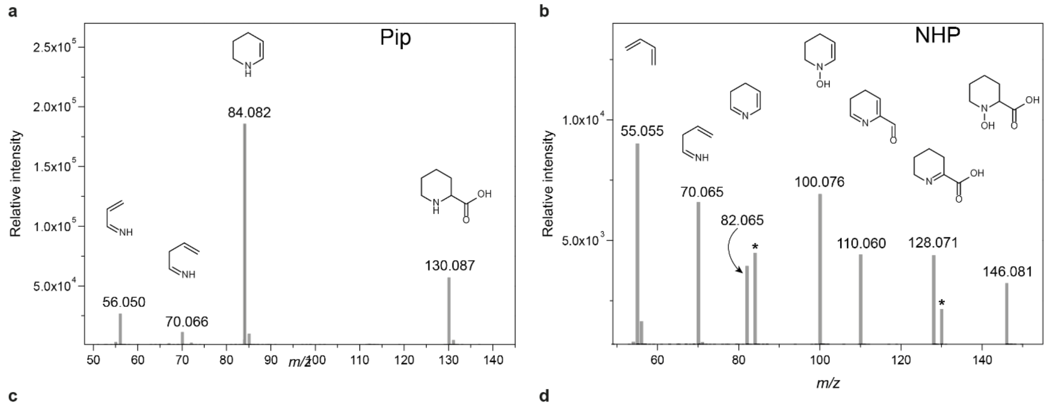


c

| MS² fragment of Pip [*m/z*] | Elemental composition | Identity |
|---|---|---|
| 130.087 | $C_6H_{12}NO_2$ | $[M+H]^+$ |
| 84.082 | $C_5H_{10}N$ | $[M-COOH]^+$ |
| 70.066 | $C_4H_8N$ | $[M-COOH-CH_2]^+$ |
| 56.050 | $C_3H_6N$ | $[M-COOH-2CH_2]^+$ |

d

| MS² fragment of NHP [*m/z*] | Elemental composition | Identity |
|---|---|---|
| 146.081 | $C_6H_{12}NO_3$ | $[M-Glc+H_2O+H]^+$ |
| 128.071 | $C_6H_{10}NO_2$ | $[M-Glc+H]^+$ |
| 110.060 | $C_6H_8NO$ | $[M-Glc-H_2O+H]^+$ |
| 100.076 | $C_5H_{10}NO$ | $[M-Glc-COOH+H_2O]^+$ |
| 82.065 | $C_5H_8N$ | $[M-Glc-COOH]^+$ |
| 70.065 | $C_4H_8N$ | $[M-Glc-C_2OOH]^+$ |
| 55.055 | $C_4H_7$ | $[M-Glc-C_2OOH-NH]^+$ |

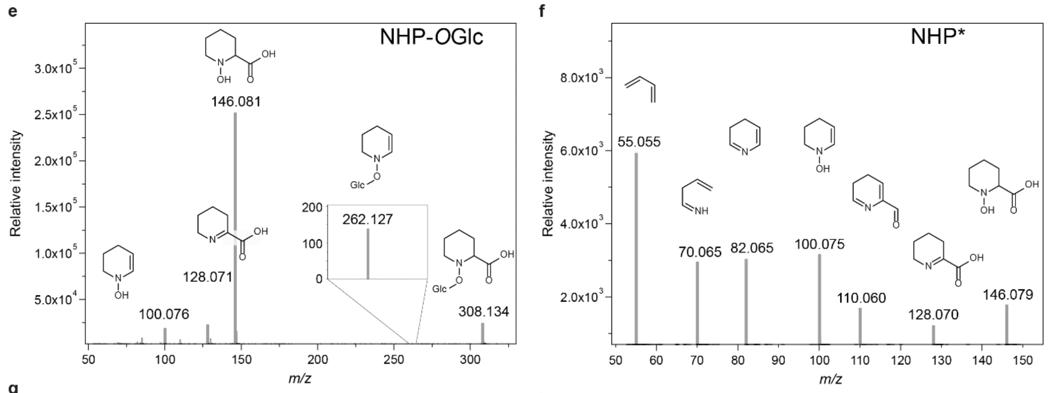


g

| MS² fragment of NHP-*O*Glc [*m/z*] | Elemental composition | Identity |
|---|---|---|
| 308.134 | $C_{12}H_{22}NO_8$ | $[M+H]^+$ |
| 262.127 | $C_{11}H_{20}NO_6$ | $[M-COOH]^+$ |
| 146.081 | $C_6H_{12}NO_3$ | $[M-Glc+H_2O+H]^+$ |
| 128.071 | $C_6H_{10}NO_2$ | $[M-Glc+H]^+$ |
| 100.076 | $C_5H_{10}NO$ | $[M-Glc-COOH+H_2O]^+$ |

h

| Pseudo MS³ fragment of NHP-*O*Glc [*m/z*] | Elemental composition | Identity |
|---|---|---|
| 146.081 | $C_6H_{12}NO_3$ | $[M-Glc+H_2O+H]^+$ |
| 128.071 | $C_6H_{10}NO_2$ | $[M-Glc+H]^+$ |
| 110.060 | $C_6H_8NO$ | $[M-Glc-H_2O+H]^+$ |
| 100.076 | $C_5H_{10}NO$ | $[M-Glc-COOH+H_2O]^+$ |
| 82.065 | $C_5H_8N$ | $[M-Glc-COOH]^+$ |
| 70.065 | $C_4H_8N$ | $[M-Glc-C_2OOH]^+$ |
| 55.055 | $C_4H_7$ | $[M-Glc-C_2OOH-NH]^+$ |

**Fig. S1. Unambiguous identification of Pip, NHP and NHP-*O*Glc by high resolution MS/MS fragmentation.** High resolution MS/MS experiments were carried out on the same samples as shown in **Fig. 1**. a **and b** MS/MS spectra of pipecolic acid (Pip) **a,** and *N*-hydroxy pipecolic acid (NHP) **b,** The fragments marked with a * in the NHP spectrum are fragments derived from glutamic acid, which elutes at a similar retention time and was selected by the first quadrupole, too. **e-f,** High resolution MS/MS **(e)** and pseudo MS³ **(f)** spectra of *N*-hydroxy pipecolic acid glycoside (NHP-*O*Glc). NHP-*O*Glc fragments during the ionization process to its aglycon. This insource derived aglycon ($[M+H]^+$ 146.081) was selected for a subsequent fragmentation to verify the identity of NHP-*O*Glc. The pseudo MS³ spectrum depicted in **f** unequivocally identifies the aglycon as NHP. The fragment annotation **(c**, **d**, **g**, **h)** is based on accurate mass information.

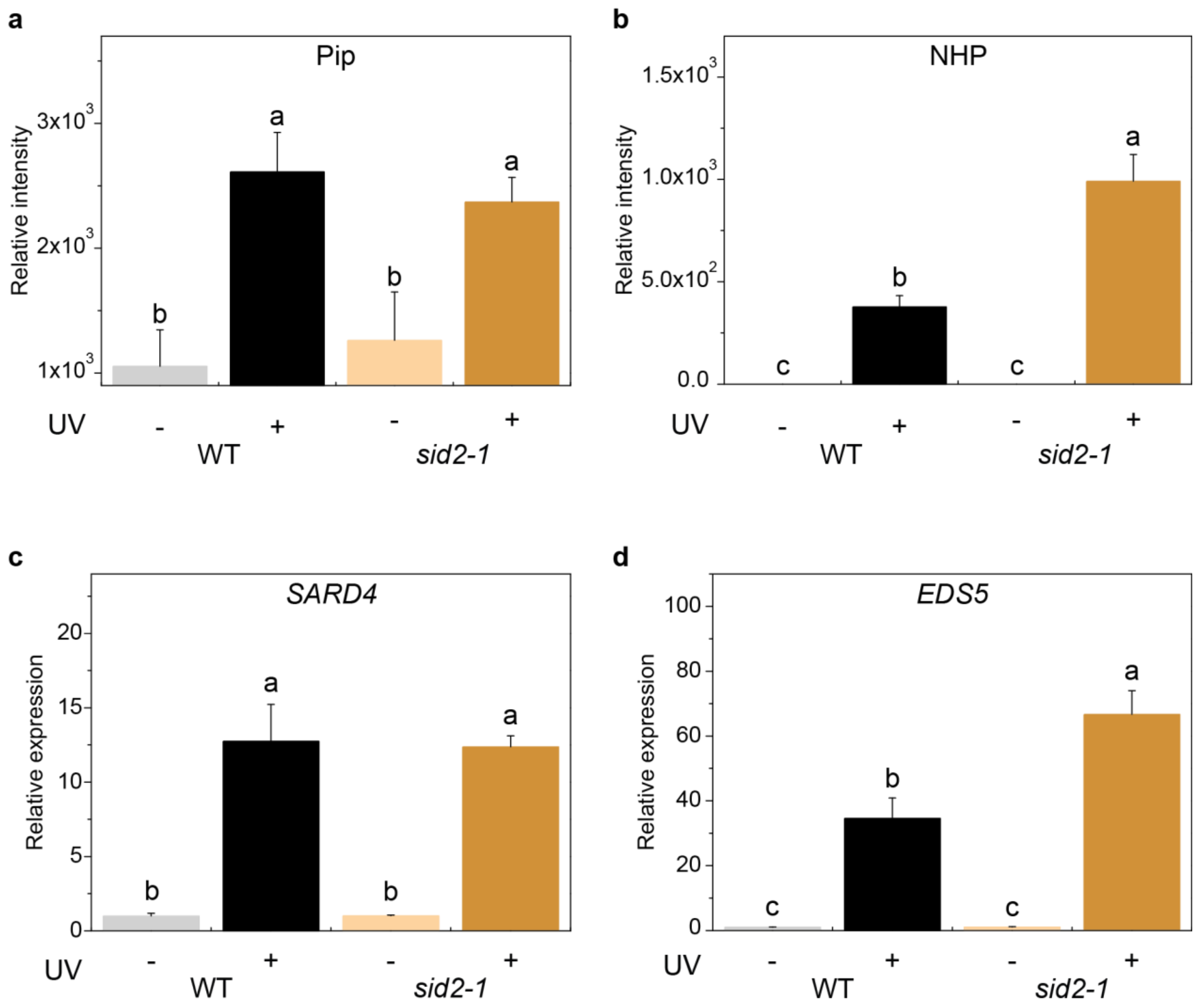


**Fig. S2. UV-C treatment leads to an accumulation of Pip and NHP, and induces the gene expression of *SARD4* and *EDS5* in the SA deficient mutant line *sid2*. a-b,** The SA deficient *sid2-1* mutant plants accumulate wild type like levels of pipecolic acid (Pip) and even more *N*-hydroxy pipecolic acid (NHP) than the wild type plants upon UV-C treatment. Samples were collected as described in **Fig. 1**. Bars represent the mean ± STD of three biological replicates. Statistical differences among replicates are labeled with different letters ($P < 0.05$, one-way ANOVA and post hoc Tukey's Test; $n = 3$). **c-d,** Relative expression levels of the Pip biosynthesis gene**,** *SARD4* (At5g52810) and the plastidial exporter *EDS5* (AT4G39030) upon UV-C stress in wild type and *sid2-1* mutant plants. Bars represent the mean ± SE of three biological replicates. Statistical differences among replicates are labeled with different letters ($P < 0.05$, one-way ANOVA and post hoc Tukey's Test; $n = 3$). The fold change of each value was normalized to the value of the untreated wild type plants.

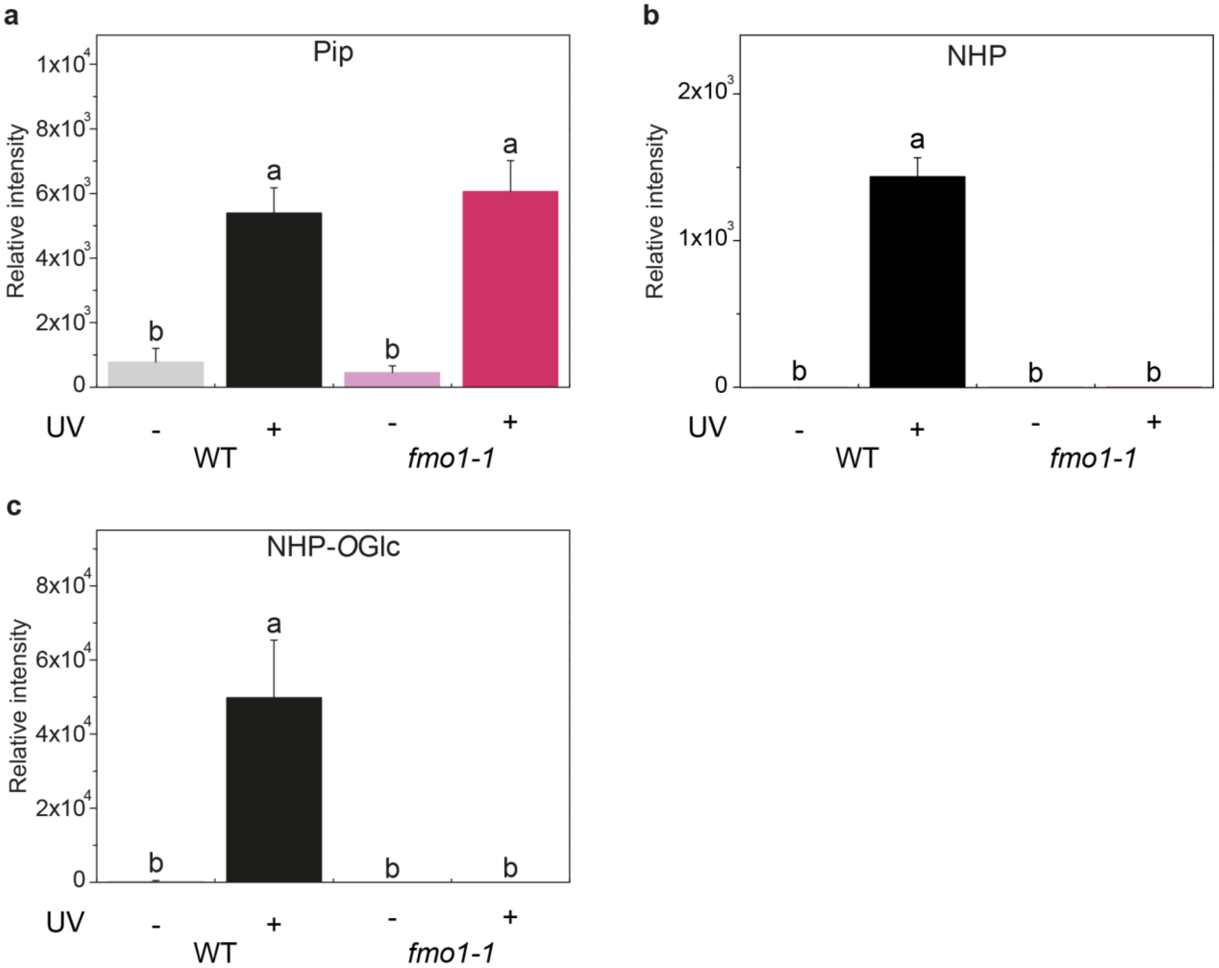


**Fig. S3. UV-C induced NHP and NHP-*O*Glc biosynthesis is abolished in *fmo1* mutant plants.** Accumulation of pipecolic acid (Pip), *N*-hydroxy pipecolic acid (NHP) and *N*-hydroxy pipecolic acid glycoside (NHP-*O*Glc) in wild type and *fmo1* mutant plants upon UV-C treatment. Samples were collected as described in **Fig. 1**. Bars represent the mean ± STD of three biological replicates. Statistical differences among replicates are labeled with different letters ($P < 0.05$, one-way ANOVA and post hoc Tukey's Test; $n = 3$). The experiment was repeated twice with similar results.

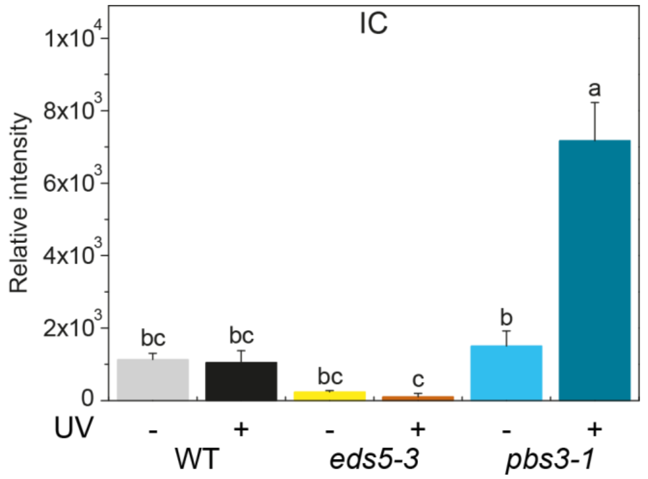


**Fig. S4. UV-C treatment induced IC accumulation in *pbs3* mutant plants, related to Fig. 1.** Samples of wild type, *eds5-3* and *pbs3-1* were collected as described in **Fig. 1**. Bars represent the mean ± STD of three biological replicates. Statistical differences among replicates are labeled with different letters ($P < 0.05$, one-way ANOVA and post hoc Tukey's Test; $n = 3$). The identity of IC was confirmed by high resolution MS/MS experiments. The experiment was repeated twice with similar results.
